# In Silico Analysis of Novel *VHL* Germline Mutations in Iranian RCH Patients

**DOI:** 10.1101/2022.10.26.513811

**Authors:** Masood Naseripour, Kowsar Bagherzadeh, Golnaz Khakpoor, Ahad Sedaghat, Reza Mirshahi, Hengameh Kasraee, Reza Afshar Kiaee, Fatemeh Azimi

## Abstract

Von Hippel-Lindau (VHL) syndrome is an autosomal dominant inherited multisystem neoplasia disorder caused by the *VHL* tumor suppressor gene, coding for VHL protein (pVHL), variants. Various types of *VHL* variants present different clinical phenotypes that later lead to events resulting in benign or malignant lesions including Retinal Capillary Hemangioblastoma (RCH). We reported on 3 novel mutation sites observed in 3 families (5 RCH patients), including c.511A>C, c.514C>T, and c.511A>T in exon 3 of the *VHL* gene. According to the ACMG classifications, c.514C>T and c.511A>T variations are likely pathogenic, and c.511A>C is a variant of uncertain significance (VUS) and in accordance with autosomal dominant inheritance. The location and impact of the incidence mutations on pVHL were computed using in silico analysis. The obtained structural information and computational analysis showed that the studied mutations induce conformational changes that limit the flexibility of the pVHL interaction interface with elonginB/C, elongin C/B, and cullin2, which is necessary for HIF1 alpha binding. The recently added gene variants and their related clinical phenotypes will improve the VHL diagnosis accuracy and the patients’ population carrying *VHL* gene mutations. These pioneering results could be a model for future functional studies.

## Introduction

Von Hippel-Lindau (VHL) syndrome is an autosomal dominant hereditary neoplastic disorder that is clinically determined by the central nervous system hemangioblastoma (CHB) and retinal capillary hemangioblastoma (RCH), renal cell carcinoma (RCC), pheochromocytoma (PCC) and other multiple tumors or cysts of the kidney (KC), liver (LC), pancreas (PC) and epididymis (EC) [1–3]. VHL syndrome can be classified as type 1 or type 2 according to the presence or absence of PCC [4, 5].

RCH is one of the most common clinical manifestations of VHL which is diagnosed approximately at age 25. It may present with progressive decreased vision which can be associated with flashing and floater [6]. More than 500 different pathogenic mutations have been recognized and studied since the detection of the VHL gene in 1993 [3–7]. Protein VHL (pVHL) forms a VCB-CR complex, including elongin C/B, cullin2, and along with RBX1/ROC1, regulates proteins ubiquitination involved in hypoxic gene response pathway [8]. Most mutations are located in two regions of high functional importance in the pVHL, including the HIF1 alpha binding site (amino acid residues 91–113) and elongin C binding site (amino acid residues 157– 171) [9].

So far five different binding surfaces (A to E) have been reported in pVHL, from which surfaces A to C are involved in the occurrence of RCH [10, 11]. Recent advancements in genetic testing technology have introduced gene mutations as standard in VHL identification and are necessary to diagnose VHL disease, especially in patients with defined implied or partial symptoms and no identified family history [12].

Advances in computational biology analysis have enabled the study of a mutation’s impact on the target macromolecules’ overall structures and monitoring of the desired structural properties without the need for time-consuming and costly experimental efforts. In this study, we described 3 families with multiple members affected with VHL syndrome based on RCH that presented at our clinic with reduced vision. In silico prediction methods and molecular dynamics simulations were employed to evaluate whether the novel variants are disease neutral or causing.

## Materials and methods

### Patients

In this case series study, 3 families (5 RCH patients) with at least one individual diagnosed with RCH were included for the *VHL* genetic testing. The employed methodologies are fully described in our previous study [13]. The resulting sequences were then submitted to the Gene Bank database (Accession numbers: ON506937, ON506938, ON506939).

#### Family A

The proband was a 31 years old woman who presented in 2018 with grade 1 fatty liver and right eye blurred vision. On examination, her pulse rate and blood pressure were 100/min and 110/75 mmHg, respectively. Her other family members have remained asymptomatic (Fig 1, A).

**Figure 1:**
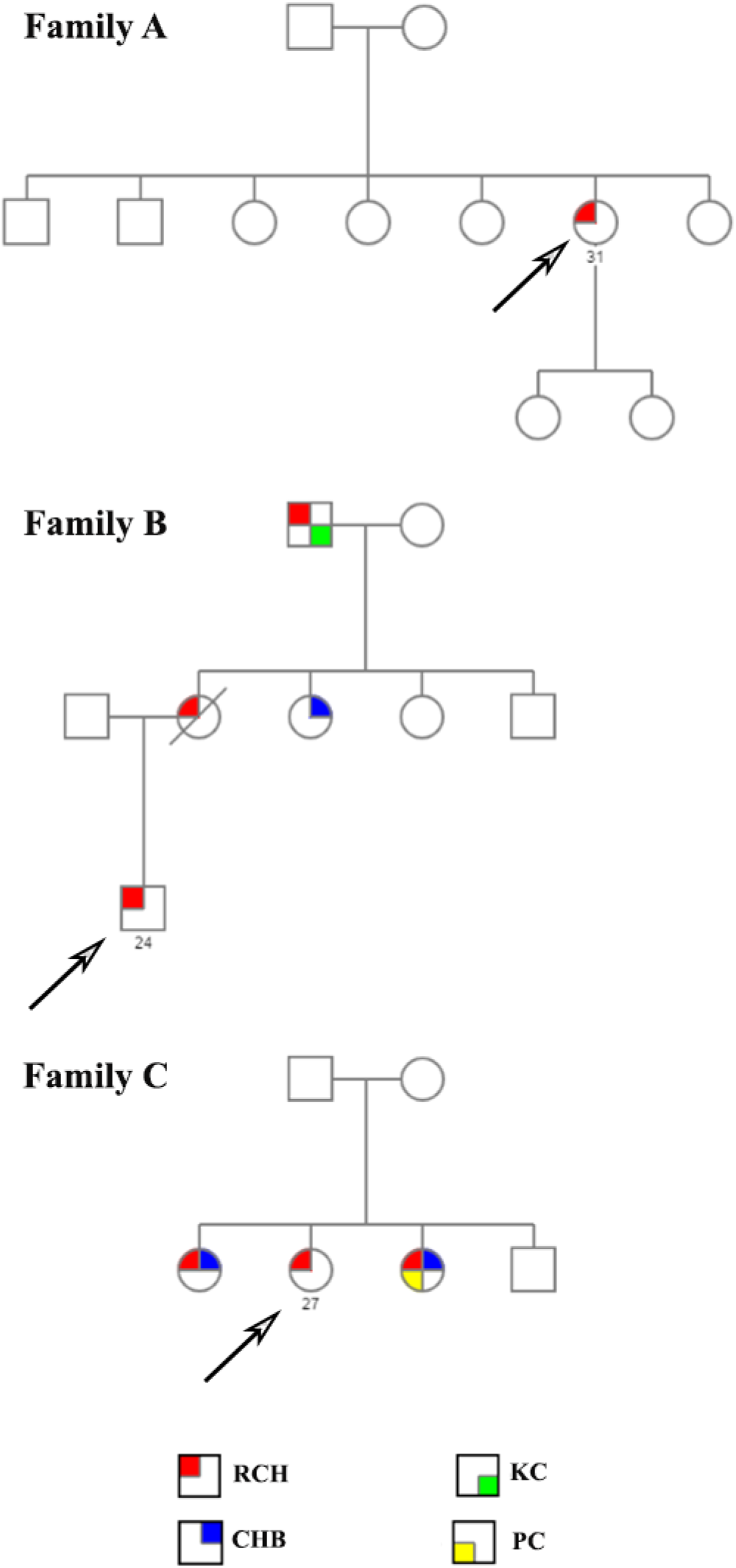
Pedigree of the affected families.

#### Family B

The proband was a 24 years old man who presented in 2020 with a right kidney cyst of 16 mm and headache. On examination, his pulse rate and blood pressure were 105/min and 135/90 mmHg, respectively. His family history was positive for RCH in his mother, CHB in his aunt, and both RCH and KC in his grandfather.

(Fig 1, B).

#### Family C

The proband was a 27 years old woman who presented in 2016 with palpitations, chest pain, and blurring of vision. On examination, her pulse rate and blood pressure were 100/min and 115/85 mmHg, respectively. Her parents have no manifestation of VHL syndrome; however, her 31 years old elder sister presented with blurring of vision, and her 21 years old younger sister with severe hypertension and blurring of vision (Fig 1, C).

### In-Silico Methodology

#### Variant Identification

The variants were first checked for new information in the Human Gene Mutation Database (HGMD). The ClinVar database (https://www.ncbi.nlm.nih.gov/clinvar/) was used to search for known variants and their clinical significance. Then the pathogenicity impact of the identified variant was discriminated employing MutationTaster (http://www.mutationtaster.org/), FATHMM-XF (http://fathmm.biocompute.org.uk/fathmm-xf/), Sorting Intolerant from Tolerant (SIFT) (https://sift.bii.a-star.edu.sg/), MutPred(http://mmb.irbbarcelona.org/PMut), PROVEAN (http://provean.jcvi.org/index.php), Varsome analysis (https://doi.org/10.1093/bioinformatics/bty897), and CADD (https://cadd.gs.washington.edu/). Deleterious thresholds were set as follows: FATHMM > 0.5, SIFT < 0.05, PMut > 0.5, and PROVEAN < − 2.5. For CADD (GRCh37-v1.6), we used the highest phred-like score cutoff recommended by the authors [14].

### Design and Simulations Procedure of pVHL Variants

The X-ray crystal structure of pVHL (PDB ID: 1lm8) was obtained from the protein data bank (https://www.rcsb.org). The initial mutant structures were retrieved by the individual substituent of the target residues to their correspondence variant using the BOVIA Discovery studio program.

Molecular dynamic (MD) simulations were carried out using GROMACS v5.1.5 software with a GROMOS AMBER force field (amber99sb-ildn) on the individual native and mutants [15, 16]. First, the models were solvated in a dodecahedral box of TIP3P water molecules [17] with a minimum distance of 14 Å between the protein surface and the box wall. Next, the net charge of the system was neutralized by replacing water molecules with appropriate counter sodium and chloride ions. The van der Waals cutoff was set to 14 Å. Periodic boundary conditions were assigned in all directions. The solvated system was then minimized through the steepest descent algorithm [18] with 1000 KJ mol-1 nm-1 tolerance followed by a canonical ensemble (NVT) and isothermal-isobaric ensemble (NPT) for 100 ps. The temperature and pressure of the system were independently maintained using the Berendsen thermostat and Parrinello-Rahman barostat algorithm at a constant temperature and pressure of 303 K and 1 bar, respectively [19]. After that, the particle mesh Ewald (PME) algorithm was employed to calculate the long-range electrostatic interactions [20]. The LINCS algorithm was applied to restrain all the bonds with an integration step of 1 fs [21]. The whole system was heated up gradually, and finally, the whole system was subjected to 100 ns of MDs at constant pressure and temperature. The stability of the computed structures was investigated by calculating the root mean square deviations (RMSD) and root mean square fluctuations (RMSF) during the simulation. The coordinate files were finally extracted from the trajectories for further analysis.

### Data Analysis and Graphical Presentation

The MDs results were analyzed and visualized with VMD 1.9 and Pymol packages [22, 23]. The graphs were all visually represented using Microsoft Office Excel [23, 24].

## Results

This study is the first report of novel heterozygous single nucleotide base substitutions in 5 RCH patients. To the best of our knowledge, the amino acid substitutions at position 171 (lysine to stop codon and lysine to glutamine) and 172 (proline to serine) have not been previously reported in VHL patients. The sequencing results from Chromas software are shown in *Supplementary Information, S1 Fig*. Given the novelty of the observed cases, an in-silico analysis was conducted to monitor the alternations at the molecular level. The results of the two-dimensional in silico characterizations of the native and mutant variants are presented in Table 1. Accordingly, the target variants were categorized as inactivating (stop mutations) and non-inactivating (missense mutations).

**Table 1:**
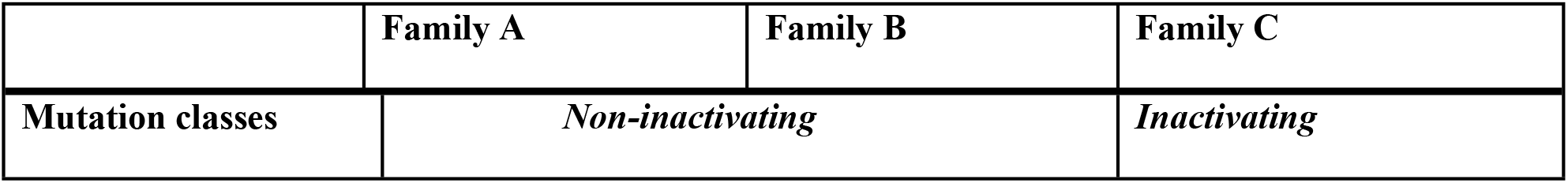
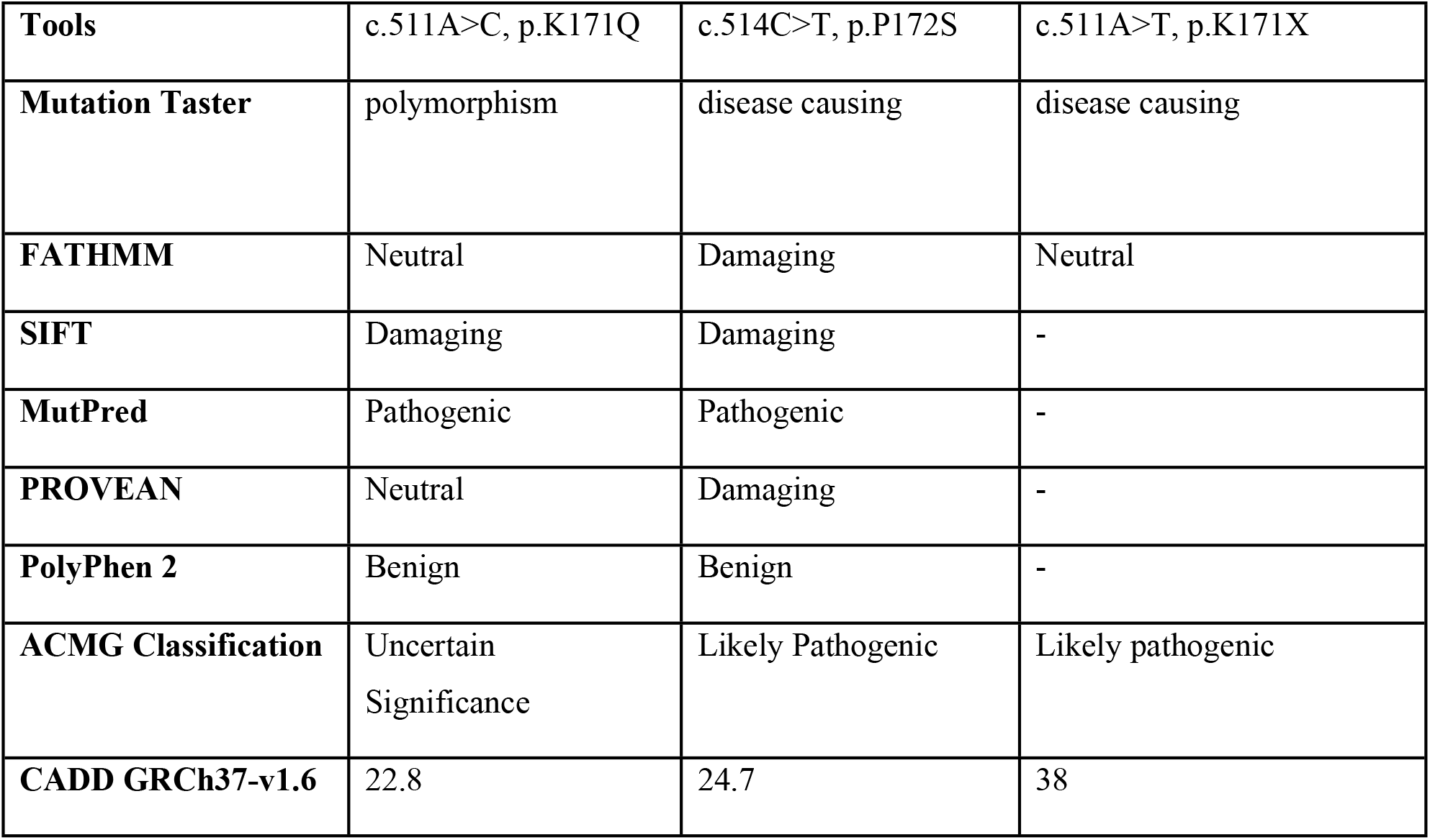
In silico prediction of exonic variants observed in the VHL gene in the Iranian RCH patients

|  | Family A | Family B | Family C |
| --- | --- | --- | --- |
| Mutation classes | *Non-inactivating* |  | *Inactivating* |
| **Tools** | c.511A>C, p.K171Q | c.514C>T, p.P172S | c.511A>T, p.K171X |
| **Mutation Taster** | polymorphism | disease causing | disease causing |
| **FATHMM** | Neutral | Damaging | Neutral |
| **SIFT** | Damaging | Damaging | - |
| **MutPred** | Pathogenic | Pathogenic | - |
| **PROVEAN** | Neutral | Damaging | - |
| **PolyPhen 2** | Benign | Benign | - |
| **ACMG Classification** | Uncertain<br>Significance | Likely Pathogenic | Likely pathogenic |
| **CADD GRCh37-v1.6** | 22.8 | 24.7 | 38 |

### Non-Inactivating Mutations

Variant c.511A>C [ENST00000256474] with genomic coordinate chr3:10191518 [*hg38 build*] leading to substitution of amino acid lysine to glutamine at position 171 [p.L171Q] in surface A, elongin B/C binding domain of pVHL (Family A) (Fig 2). This mutation was classified based on the ACMG as VUS. Clinical manifestations of c.511A>C and c.514C>T variants had only RCH. After testing other family members, we found that c.511A>C variant occurs sporadically without systemic involvement.

**Figure 2.**
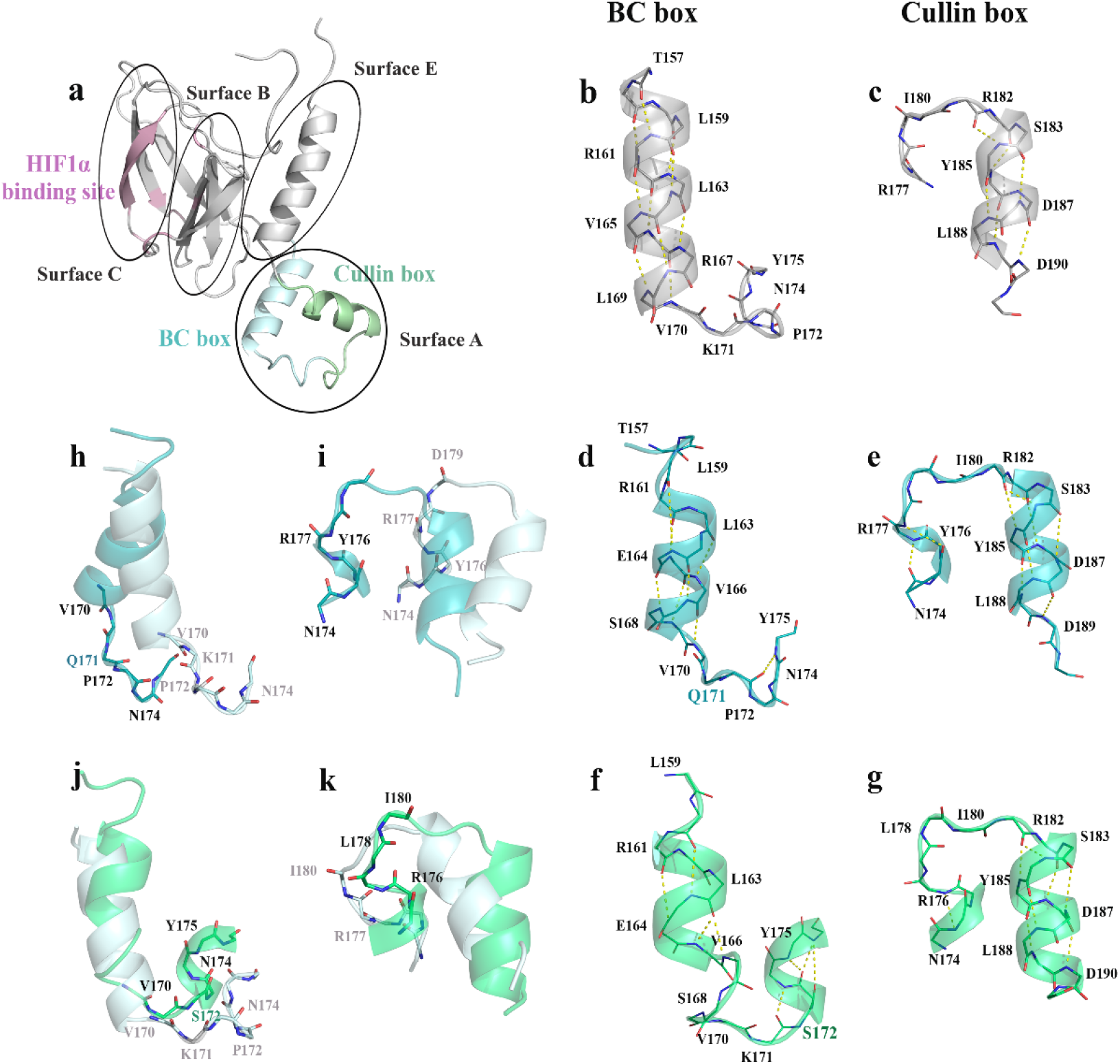
**a**. The overall structure and regions of native *pVHL*, **b**. The amino acid arrangement in BC box located in *pVHL* α-domain, **c**. The amino acid arrangement in cullin box, located in *pVHL* α-domain, **d**. The amino acid arrangement in BC box located in mutant K171Q α-domain, **e**. The amino acid arrangement in cullin box, located in mutant K171Q α-domain, **f**. The amino acid arrangement in BC box located in mutant P172S α-domain, **g**. The amino acid arrangement in cullin box, located in mutant P172S α-domain, **h**. Superimposition of the amino acid in native *pVHL* BC box over the correspondence residues in mutant K171Q α-domain, **i**. Superimposition of the amino acid in native *pVHL* cullin box over the correspondence residues in mutant K171Q α-domain, **j**. Superimposition of the amino acid in native *pVHL* BC box over the correspondence residues in mutant P172S α-domain, **k**. Superimposition of the amino acid in native *pVHL* cullin box over the correspondence residues in mutant P172S α-domain.

The other variant of c.514C>T, with genomic coordinate chr3:10191521 [*hg38 build*], that leads to substitution of amino acid proline to serine at position 172 [p.P172S] is also located in surface A, the in the Eelongin B/C binding domain of pVHL (Family B). However, this mutation is classified as likely pathogenic. While the clinical manifestation of this variant had RCH, other family members were not accessible for evaluation. After a detailed literature and database review, no evidence was found that, the target variants have been previously reported as a type of VHL causative.

### Inactivating Mutation

The nonsense mutation c.511A>T [ENST00000256474, NM_000551] with genomic coordinate chr3:10191518 [*hg38 build*] generated premature termination codon (PTC) at position 171[p.K171X] to substitute amino acid lysine to stop codon at in elongin B/C binding domain of pVHL (Family C). This mutation was also observed in 2 probands sisters with a likely *VHL* type 1 phenotype. However, the mutation was not detected in her parents or brother. Therefore, the mutation is classified as likely pathogenic. Clinical presentations of the mutation varied from isolated RCH in the proband to concurrent presentation of CHB and PC in the proband’s sisters.

### In-Silico Characterization of the Studies pVHL Native vs. Mutant Structures

The studied structures were submitted to a comprehensive set of atomistic MDs to compute the impact of the target mutations on the pVHL structure compared with that of the native form. Basic MDs trajectory analysis was performed, including root mean square deviations (RMSD) and root mean square fluctuation (RMSF), as the time evolution of target conformations using GROMACS software.

The backbone RMSD plots of the studied structures showed that the trajectories become stable after 60 ns, 75 ns, 35 ns, and 10 ns of MD simulations for the native, K171Q, P172S, and K171X structures (Supplementary Information, S2 Fig). Accordingly, the root means square deviations plot of the K171X structure shows that the omission of surfaces A and E dramatically decrease the deviations due to the presence of the rigid antiparallel β-sheets in surfaces B and C. Furthermore, K171Q mutation stabilizes the overall protein structure while P172S mutations induce a structural rearrangement (like refolding of a loop into α-helix or vice versa), which positions a higher percentage of residues in this mutant’s structure compared to the native and the one carrying the P172S mutation (Supplementary file, S1Table).

The per-residue root-mean-square fluctuations (RMSF) of the native and the studied mutants were also computed (Supplementary file, S2 Fig). As observed, the root means square RMSF in the three mutants have significantly decreased in comparison with the native structure. The number of hydrogen bond formed in the overall studied structures as well as the surface accessible areas was computed and graphed (Supplementary Informations, S 3 – 4 Fig).

## Discussion

The variants identified in our study are all heterozygous and are inherited autosomal dominant. The final diagnosis and subsequent clinical follow-up depend on its probable likely pathogenic interpretation, which is an important step. This interpretation is straightforward if it involves destroying the protein function by creating a premature termination codon (PTC) or changing the transcript; otherwise, it will not be easy to interpret [25].

In the current study, the effect of each novel variant and the evaluated impact of mutated amino acids on protein folding and stability were investigated with the aim of predicting pathogenic molecular mechanisms in VHL disease.

Web-based programs were employed to estimate whether the amino acid substitutions in our study can affect protein function, including Mutation Taster, FATHMM-XF, SIF, Mut Pred, PROVEAN, Varsome analysis, and CADD. ACMG Classification revealed 2 new variants, one Likely Pathogenic and one variant reported as VUS.

### Inactivating Mutations

It has already been shown that mutations that lead to PTC due to disruption of the α domain in terms of vital structure may have a powerful pathogenic role, which in interaction with Elongin C / B, forms a 3D protein structure.^26^ The inactivating variant c.511A>T in exon 3 identified in our patients results in a lysine that stops codon substitution at amino acid 171 in the elongin C binding domain of pVHL. *In vitro* residues scanning studies have clearly displayed that residues from 157 −171 on pVHL constitute an elongin C binding site and have been a hot spot for VHL-causing mutations [8, 27]. Also, similar to Minervini et al.’s 2019 study, this variant is placed on surface A (154-189), which is related to the RCH development [11]. We predicted that the omission of surfaces A and E increase the rigidity and might have problems in an interaction with a VCB-CR complex.

The p.K171X might escape the nonsense-mediated mRNA decay (NMD) machinery because the PTC is located in the last exon of VHL, and the NMD typically eliminates mRNA transcripts containing premature stop codons and the presence of at least one intron, similar to the p.Tyr175X which described previously in other studies [28,29]. Also, if this transcript is processed, it still produces the pVHL molecule, which lacks the main part of the α-domain [25,26], showing high pathogenicity.

### Predicted Effects of Novel Missense Variants

Two novel missense variants (c.511A>C and c.514C>T) were detected in 2 unrelated families (Table 1). Of these, the proband presented with an apparently sporadic RCH, and the other proband had a clinical presentation in his family. The performed in silico studies show that 2 missense variants are located on the *α*- domain; one of these, the c.511A>C transition, is located at the elongin C binding site. Substitutions that occur in the *α*-domain (vital for protein binding) may impact interaction.

The protein structure’s overall fluctuations show a significantly lower degree of flexibility in the mutant structures compared to the native structure. The per residue fluctuation analysis reveals a considerably different pattern of fluctuations in α-domain participant amino acids in the target mutants (except for that of K171X, where the region is missed), showing that the mutation dramatically affected the flexibility of the protein. Previous structural analysis of pVHL has emphasized the necessity of flexibility in the α-domain region, which is needed for conformational change upon interactors binding [10]. The positively charged patch contributed by K171 resulted in the establishment of attractive charge interactions with E173 as well as hydrogen bond formation with N174 through the lysine hydroxyl group. Substitution of K171 with Q results in the formation of amide-π interactions between the Q171 carbamoyl group and Y175 that extends the electrostatic interactions between H1 and H3 to H2, which enhances the electrostatic interactions network between the helixes (Fig 3), limiting the participant residues fluctuations. Substitution of P172 with serine allows the region to adapt to a helical geometry through hydrogen bond formation with Y175 (Fig 2). The substitution also enhances the electrostatic interactions network between the helixes (Fig 3) as well as a hydrogen bond formation between S172 and D189 that further deplete the region flexibility.

**Figure 3.**
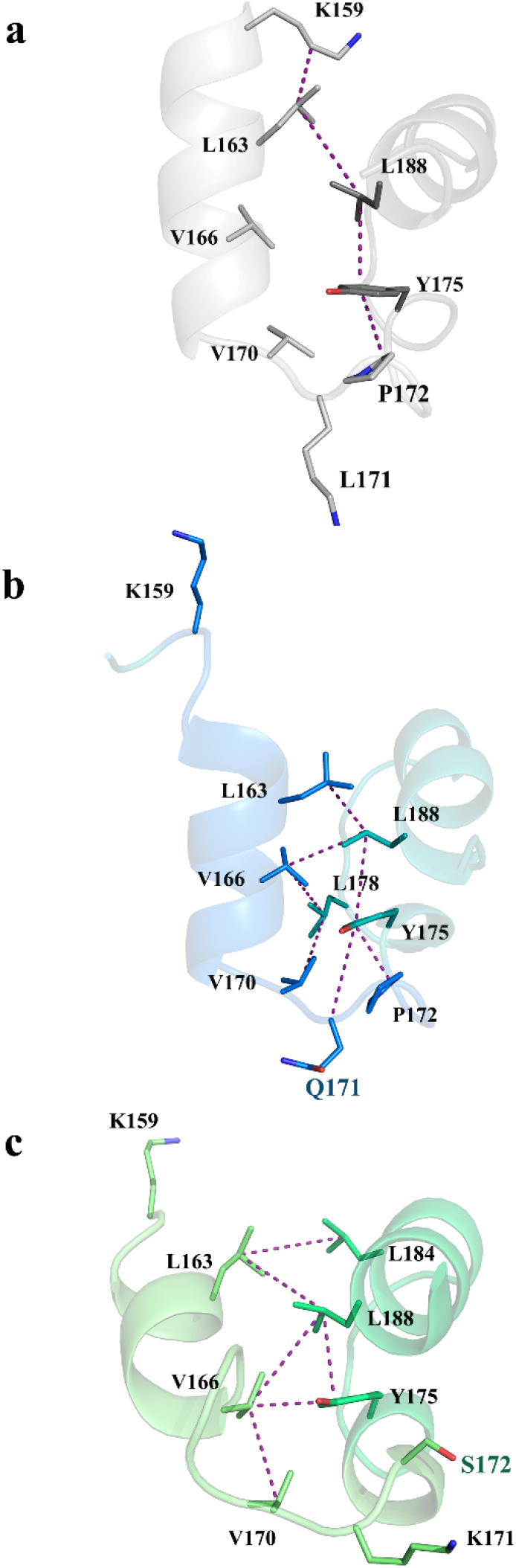
The comparison between the electrostatic interactions between H1, H2, and H3 residues interacting residues in **a**. the Native, and **b**. K171Q and **c**. P172S.

Although the target mutations result in both BC box and Cullin box refolding, the rearrangement in the BC box changes the region conformation drastically, while the induced changes resulting from P172S are more significant than that of K171Q (Fig 2). According to the obtained results, it is inferred that the observed mutations restrain the α-domain region flexibility, resulting in the enhanced stability of mutant structures RMSD plots. Likewise, no significant difference is observed in the number of amino acids formed between the constructed amino acids in the native and mutants (except for that of K171X).

As previously shown, any conformational changes in the pVHL α-domain are induced through an allosteric pathway (Fig 4) that starts from the interactions between the P564 residue of HIF-1α with residue S111 in the β-domain of pVHL and extends to S168 in the α-domain. The pathway enhances the interactions between β-domain and α-domain to restrain the α-domain flexibility, which favors elongin C binding. The impact of the observed mutations over the allosteric pathway was also computed. Accordingly, the mutations alter the localization of residues Q164 and S168 in the BC box alpha-helix, ultimately disturbing the hydrogen bond network in the pathway and affecting interactions with elongin C and, in turn, a host of other molecules [30].

**Figure 4.**
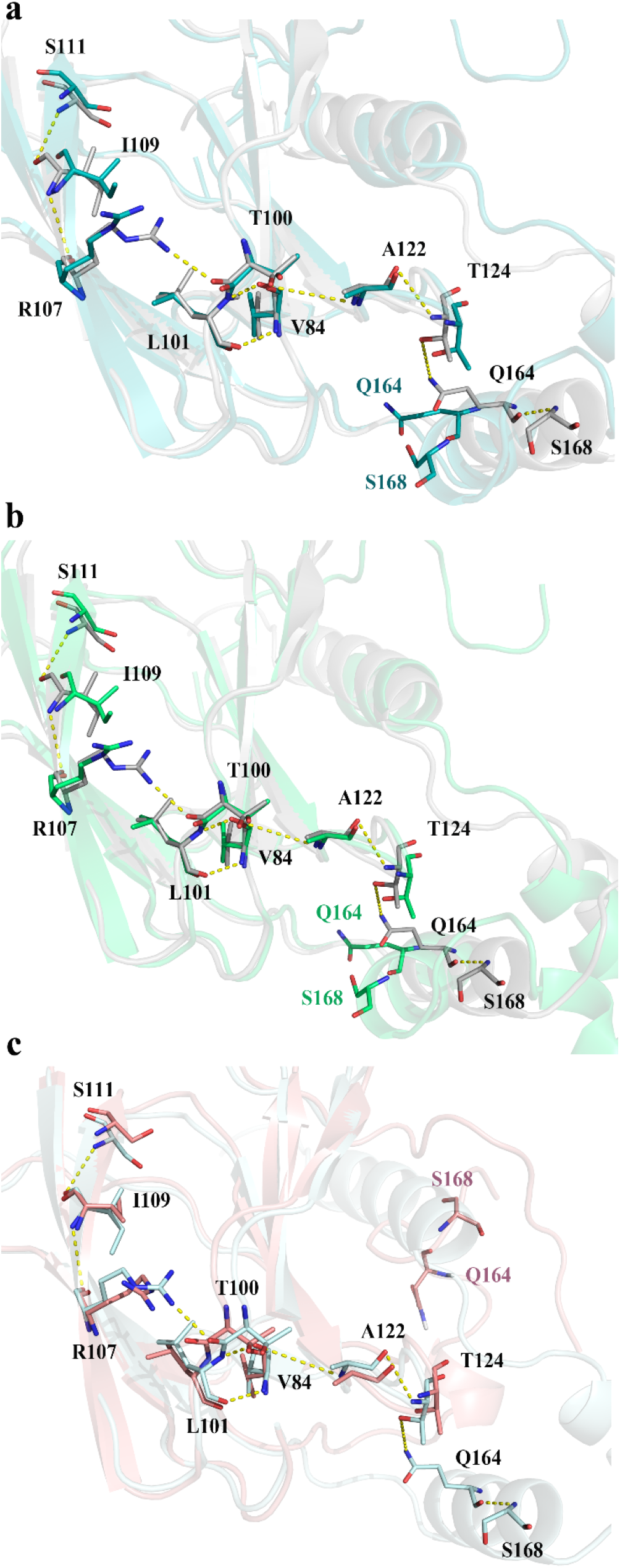
Superimposition of the allosteric pathway from S111 to S168 in α-domain native *pVHL* over the mutant correspondence, **a**. K171Q mutant structure, **b**. P172S mutant structure and, **c**. K171X structure.

Further, hydrogen bond analyses of the native and mutant proteins with respect to the simulation time were performed (Supplementary Information, S3 Fig). No significant difference was observed between the numbers of hydrogen bonds formed during the simulation (except for mutant K171X, in which the smaller number of hydrogen bonds is obviously due to the loss of the whole α-domain). Therefore, it seems that the region’s flexibility is altered more through electrostatic interactions (Fig 3). The solvent-accessible area (SASA) of the native and mutant protein trajectory values was calculated (Supplementary Information, S4 Fig). Accordingly, the SASA value is lower in mutants K171Q and P172S in comparison with the native structure. Furthermore, the mutant residues and their adjacent amino acids accessibility have decreased significantly, showing that the mutations could affect the tertiary structure of the proteins, and the mutant structures’ interactions with elongin B/C would be affected and probably not possible. In conclusion, based on the in-Silico characterizations, both K171Q and P172S mutations induce conformational rearrangement in the protein structure in such a way that the α-domain participating residues fluctuations are limited, and the rigidity increases.

### Genotype-Phenotype Correlation

Nonsense variants or exonic deletions leading to the loss of functional mutations are often associated with the VHL type 1 phenotype [31]. The subjects carrying the p.K171X variant presented a variety of manifestations from isolated RCH to Von Hippel-Lindau syndrome. It is expected to produce VHL proteins largely lacking the C-terminal region.

Moreover, missense variants in the elongin C binding site in the α- domain cause Type 1 VHL and, through the involvement of the main acid amines, lead to a complete loss of pVHL function [32]. Key amino acids caused the proband carrying the p.K171Q variant to completely lose the elongin C binding domain function of pVHL [9]. This variant only manifested isolated RCH. For example, the proband having p.P172S had only RCH, but other family members had wide ranging clinical presentations.

We showed that the c.514C>T and c.511A>T variations in the elongin C binding α-domain of pVHL might be disease-causing. Disease-causing mutations are often located in highly conserved positions in a protein sequence because of their functional importance [33]. However, it is difficult to predict the phenotype even using the entire database of mutations and basic molecular research, because of variability in systemic abnormalities is an important feature of this disease.

## Conclusions

In silico prediction strategy and results of genotype analysis of RCH patients provide potential evidence for the pathogenicity of unreported variants in exon 3. The three novel mutations evaluated in the present study have significantly decreased fluctuation in comparison with the native structure and increased rigidity. These findings will undoubtedly add new information to the VHL gene databases. Furthermore, the results of this pilot study will provide a basis for future comprehensive studies.

## Acknowledgments

The authors thank all those who provided support during this research.

## Author’s contributions

M.N: Introduced and treated the patients and revised the final; K.B. performed Molecular dynamic and 3D structure and revised the manuscript text; G.KH: Genetic consultation; A.S: Introduced the patients; R.M: wrote the clinical findings; H.K & R.A: collected clinical findings; F.A: wrote the Initial draft of the manuscript, samples collection and genetic analysis, genetic tests interpretation.

## Supporting information

In Silico Analysis of Novel VHL Germline Mutations in Iranian RCH Patients

---

**S1Table.** The participant residues in different structures of pVHL during 100ns of MDs (all values are in percent (%)).

| Variant | Sructure | Coil | β-sheet | B-bridge | Bend | Turn | A-helix | 3-helix |
| --- | --- | --- | --- | --- | --- | --- | --- | --- |
| Native | 0.54 | 0.31 | 0.24 | 0.03 | 0.12 | 0.13 | 0.15 | 0.04 |
| K171Q | 0.54 | 0.29 | 0.24 | 0.02 | 0.12 | 0.13 | 0.15 | 0.04 |
| P172S | 0.55 | 0.32 | 0.23 | 0.03 | 0.10 | 0.14 | 0.15 | 0.03 |
| Missense | 0.54 | 0.32 | 0.32 | 0.04 | 0.12 | 0.14 | 0.04 | 0.02 |

**S1 Fig.**
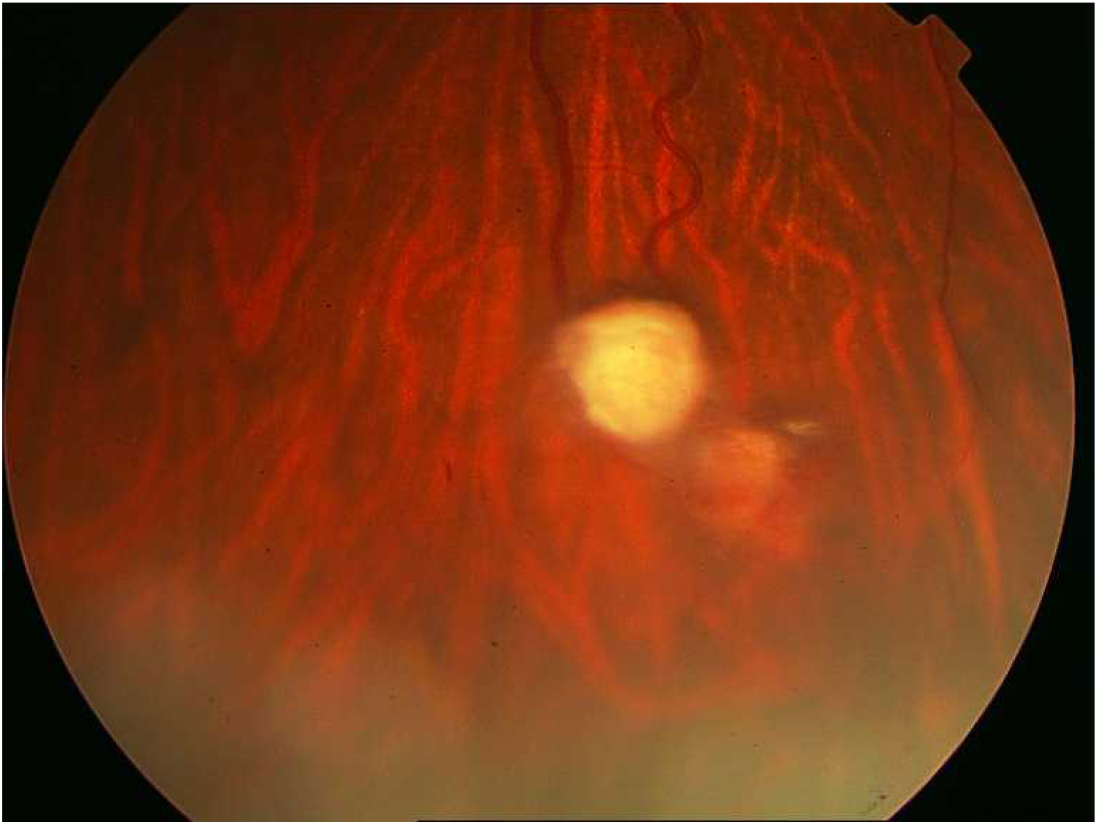
The lesions treated with direct laser photocoagulation

**S2 Fig.**
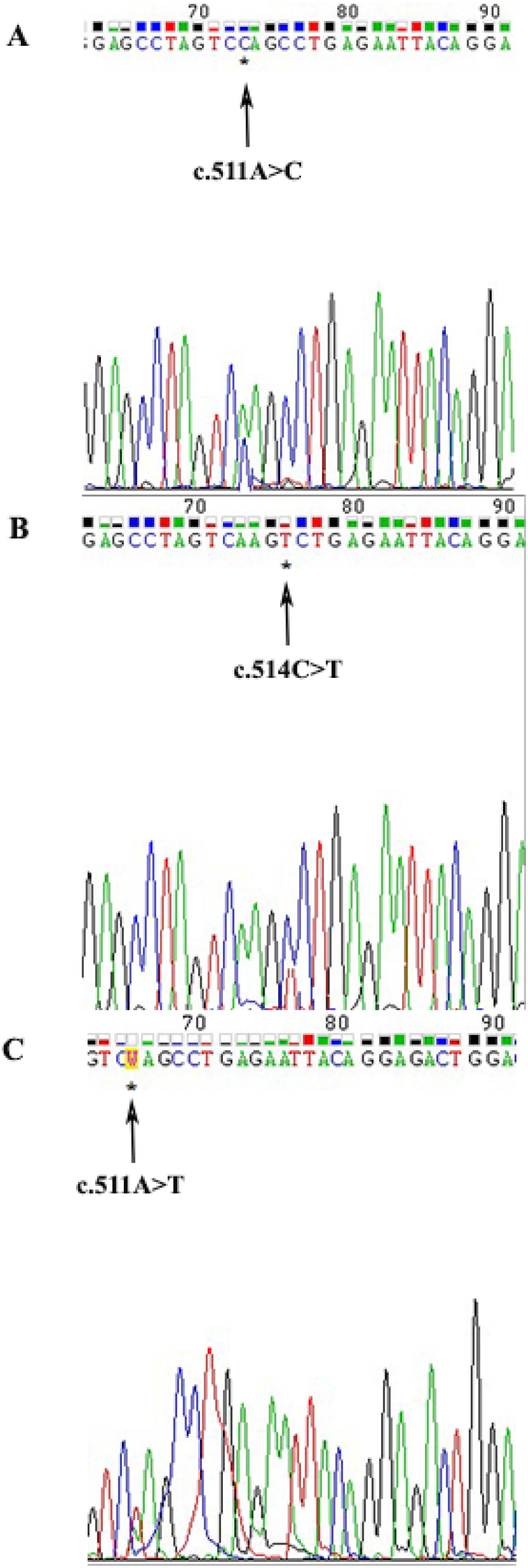
Electropherogram showing the heterozygous mutation A: c.511A>C, B: c.514C>T, and C: c.511A>T in exon 3.

**S3 Fig.**
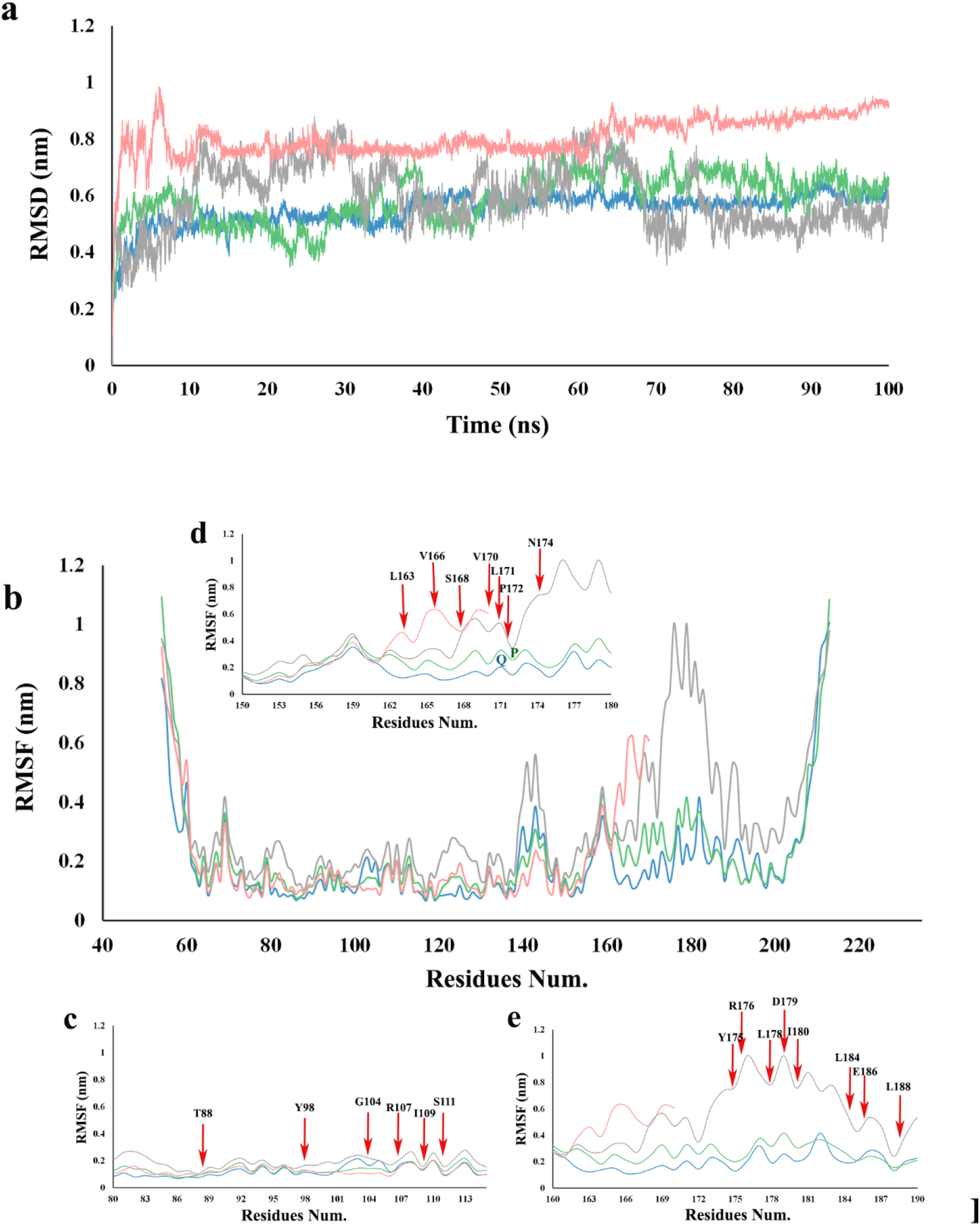
**a.** Back-bone root mean square deviations (RMSD) all the studied structures (Native (gray), K171Q (blue), P172S (green), Missense (Pink), **b**. Per residue root mean square fluctuations (RMSF) of all the studied structures (Native (gray), K171Q (blue), P172S (green), Missense (Pink), **c**. close view of RMSF plots for residues 80-113, **d**. close view of RMSF plots for residues 150-180, and **e**. close view of RMSF plots for residues 160-190.

**S4 Fig.**
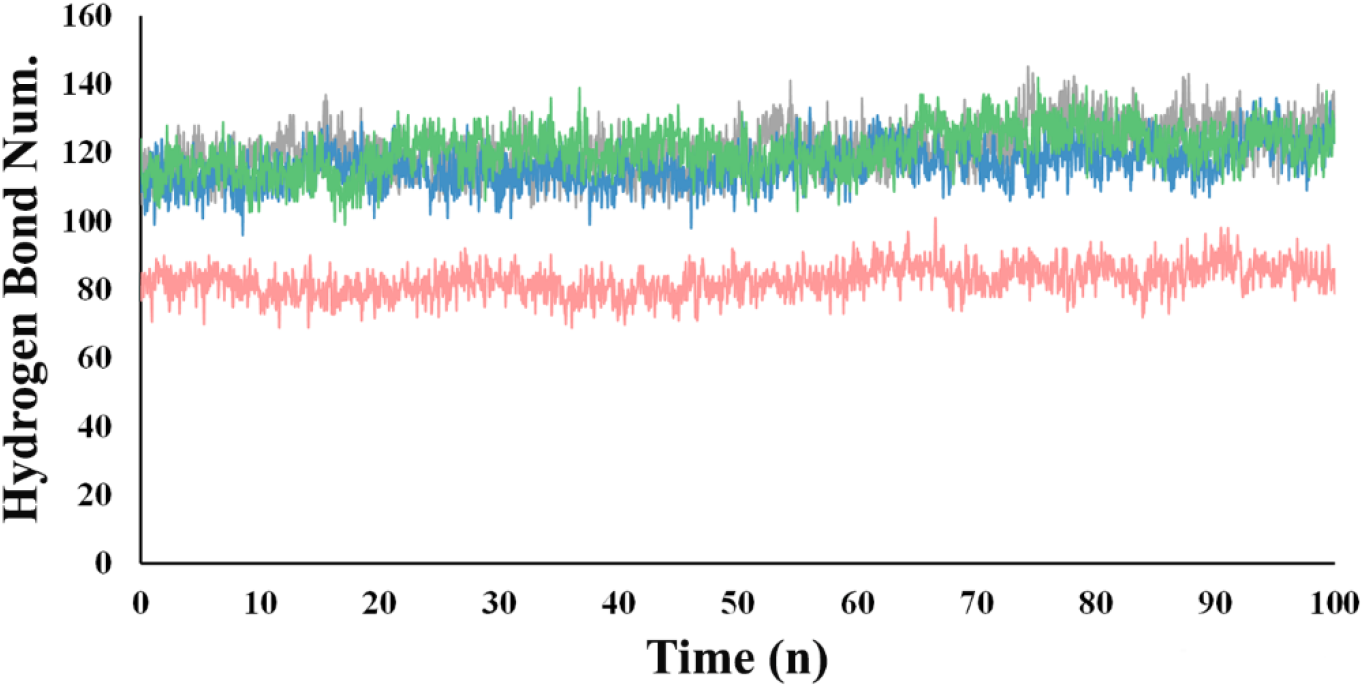
Total number of hydrogen bonds formed in the native and mutant *pVHL* structures (Native (gray), K171Q (blue), P172S (green), Missense (Pink),

**S5 Fig.**
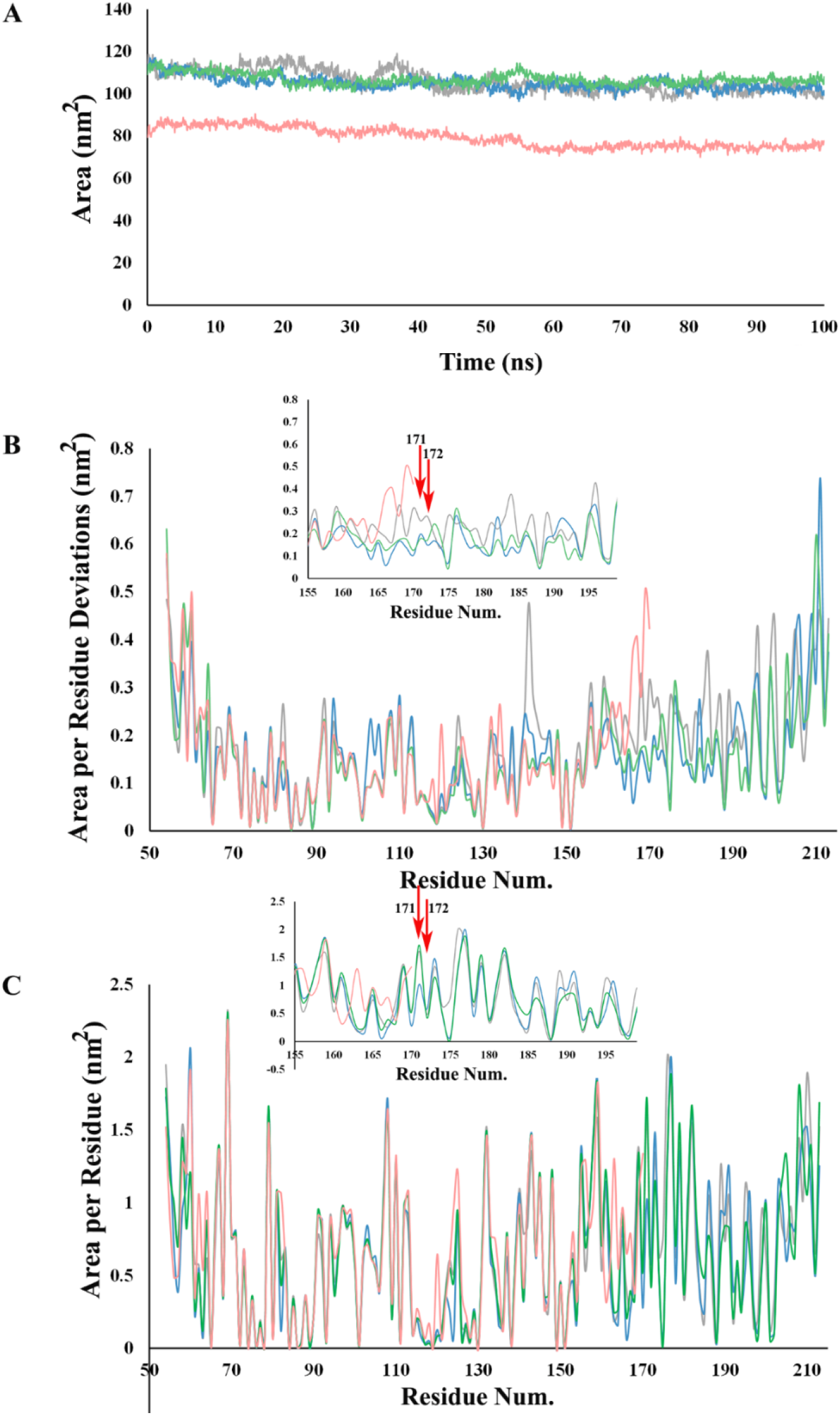
Solvent accessible surface area (SASA) analysis of native and mutant structures *pVHL* (Native (gray), K171Q (blue), P172S (green), Missense (Pink), **a**. SASA of the overall structure, **b**. SASA deviation per residue, and **c**. SASA per residue

